# Elucidating human ageing-related phenotypic abnormalities with a novel hierarchical feature selection-based knowledge discovery framework

**DOI:** 10.1101/2023.05.26.542395

**Authors:** Cen Wan, Carl Barton

## Abstract

Ageing is a complex biological process involving multiple genes that are also related to phenotypic abnormalities. However, the micro view of the associations between ageing and human phenotypic abnormalities is still under-studied. We propose a new hierarchical feature selection method namely HIP_+_, which selects positive hierarchical information-preserving features according to pre-defined hierarchical ontology information. The experimental results confirm that HIP_+_ obtained better performance than the state-of-the-art hierarchical feature selection method in predicting human phenotypic abnormalities annotations. We further propose a HIP_+_-based knowledge discovery framework that also successfully highlights some important associations between biological processes and human phenotypic abnormalities.

## 1 Introduction

Ageing is a complicated process that involves multiple biological pathways and the associated genes. Many of those genes are also closely related to human phenotypic abnormalities like global developmental delay^1^ and aplasia of the cerebrum^2^. However, in general, the micro view of the associations between ageing and human phenotypic abnormalities is still under-studied. Thanks to the development of biological databases that have already stored an increasing number of information about human genes, the study of ageing and phenotypic abnormalities now can be boosted by machine learning methods. In this work, we propose a novel hierarchical feature selection-based knowledge discovery framework to highlight some important associations between ageing and phenotypic abnormalities.

The hierarchical relationships (i.e. generalisation-specialisation or is_a relationships) are commonly used to organise multiple semantic concepts (a.k.a. terms) as a tree-like structure. The terms that are close to the root usually bear more generic definitions, whereas the terms that are close to the leaves bear more specific definitions. In this work, we use two well-known bioinformatics databases, i.e. Gene Ontology (GO)^3^ and Human Phenotypic Abnormalities (HPO)^4^. The former uses unified vocabularies (GO terms) to describe gene functions, whilst the latter defines genes’ corresponding phenotypic abnormalities encountered in human disease. Both databases use hierarchical relationships to organise a large number of biological and medical definitions. For example, the root term GO:0008150 bears the most generic definition of Biological Process, whilst a leaf term GO:0010588 defines a specific type of biological process – cotyledon vascular tissue pattern formation. Therefore, according to the hierarchical relationships, if one gene is annotated with one term, all ancestors of that term will also be annotated with that gene. But this principle also leads to a difficulty in knowledge discovery, since the dimensionality of the derived dataset is usually very high.

Hierarchical feature selection^5–11^ is an emerging topic in dimensionality reduction research. It exploits pre-defined hierarchical feature relationships to remove redundancies between features, leading to improved predictive performance. For example, the HIP method^7,12^ is one of the best-performing hierarchical feature selection methods which merely selects those features that preserve the most hierarchical information. A large-scale empirical evaluation^9^ suggested that HIP leads to the general best predictive accuracy when using Gene Ontology terms as features to predict pro/anti-longevity genes. Another important merit of the HIP method is its unsupervised learning setting – it merely conducts feature selection according to a pre-defined hierarchy, without considering samples’ label information. As inspired by recent research on hierarchy-aware machine learning methods^13–15^ – prioritising positive feature values improves predictive accuracy, we propose a novel hierarchical feature selection method namely HIP_+_ that also selects the hierarchical information preserving features bearing positive values. This paper is organised as follows. Section 2 introduces the newly proposed HIP_+_ feature selection method and knowledge discovery framework. Section 3 presents the computational experimental results and discusses some highlighted Gene Ontology terms that are selected by the HIP_+_ method. Section 4 presents conclusions and future research directions.

## 2 Methods

### 2.1 Overview of the HIP+ hierarchical feature selection method and knowledge discovery framework

We proposed a Gene Ontology-based knowledge discovery framework using, namely HIP+, to highlight the relationships between ageing-related genes and human phenotypic abnormalities. As shown in Figure 1, we created a feature matrix by using Gene Ontology terms (GO database version 2022-7-1) to predict a set of Human Phenotype Ontology terms (HPO database version 2022-6-11) for 184 human ageing-related genes reported in the GenAge database (Build 20)^16^. Then the proposed HIP_+_ method was applied to both types of ontology terms. The HIP_+_ method was first applied to the GO term matrix in order to reduce the dimensionality. The selected GO terms by HIP_+_ also bear the most specific biological definitions of individual genes. Analogously, due to the hierarchical organisation of HPO terms, the class label matrix (i.e. HPO terms) was also transformed by using the HIP_+_ method, in order to select the most specific HPO terms annotated for individual genes since more specific HPO terms’ definitions are more valuable for the knowledge discovery purpose. Note that, before applying the HIP_+_ method on the HPO terms class label matrix, we firstly filter some rare HPO terms (i.e. with very few annotated genes) by applying a threshold value *τ*, denoting the lowest percentage of annotated genes for each HPO term. Finally, after applying the HIP_+_ method on both matrices, a random forests^17,18^ classifier will be trained and used to predict the HPO terms annotation by using GO terms. The feature importance information derived by the random forests classifier is used for interpreting GO terms w.r.t. HPO terms prediction.

**Figure 1.**
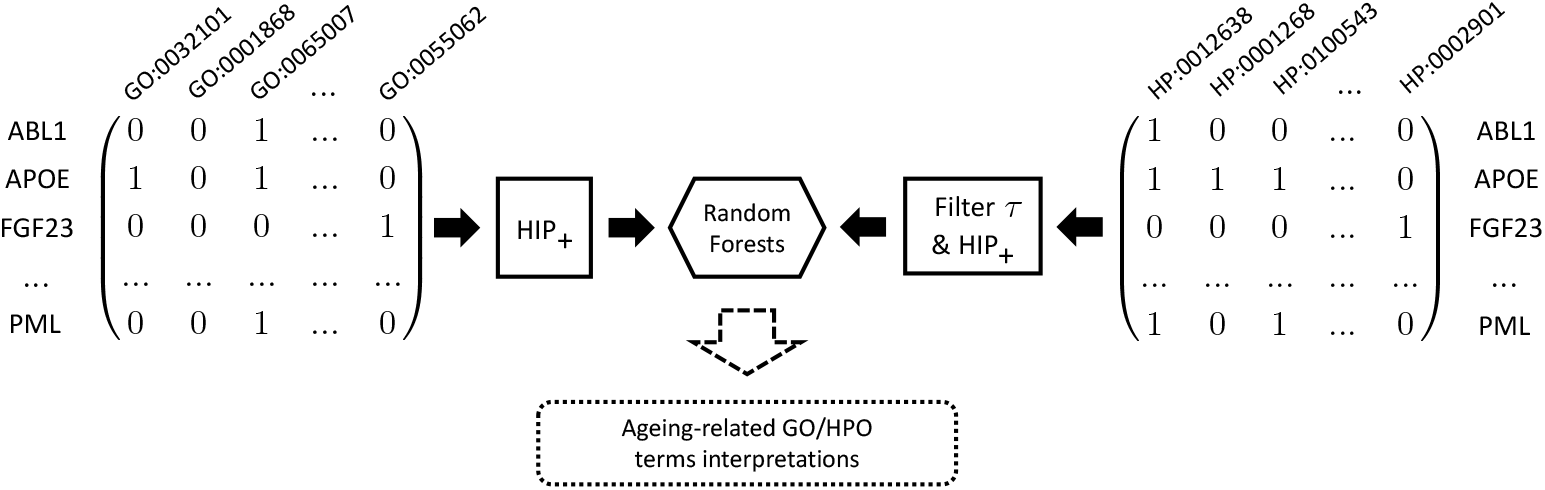
The overview of the HIP_+_ hierarchical feature selection-based knowledge discovery framework.

### 2.2 Selecting hierarchical information preserving features bearing positive values (HIP_+_)

HIP_+_ is a natural extension of the well-known HIP^7^ feature select method. It prioritises the features that bear positive values and preserve the most hierarchical information simultaneously. The pseudocode of the HIP_+_ method is shown in Figure 2.a, which takes five variables as the inputs, i.e. a set of annotated terms 𝒯_*g*_ and their values 𝒱(*g*) for the target gene *g*; the ancestor (𝒜(𝒯_*g*_) and descendent 𝒟(𝒯_*g*_) sets for each individual terms and the selection status for each individual terms. From lines 1–12, HIP_+_ firstly checks the feature values – if the feature value for one term equals *1*, then all the ancestor terms in the hierarchy will be removed from the selection term set; *vice versa*, if the value of one term equals to *0*, then that term and all its descendent terms will be removed. This selection process will be repeated for each individual term in the given dataset. From lines 14–18, all the selected terms are used to create a new dataset for the downstream tasks.

**Figure 2.**
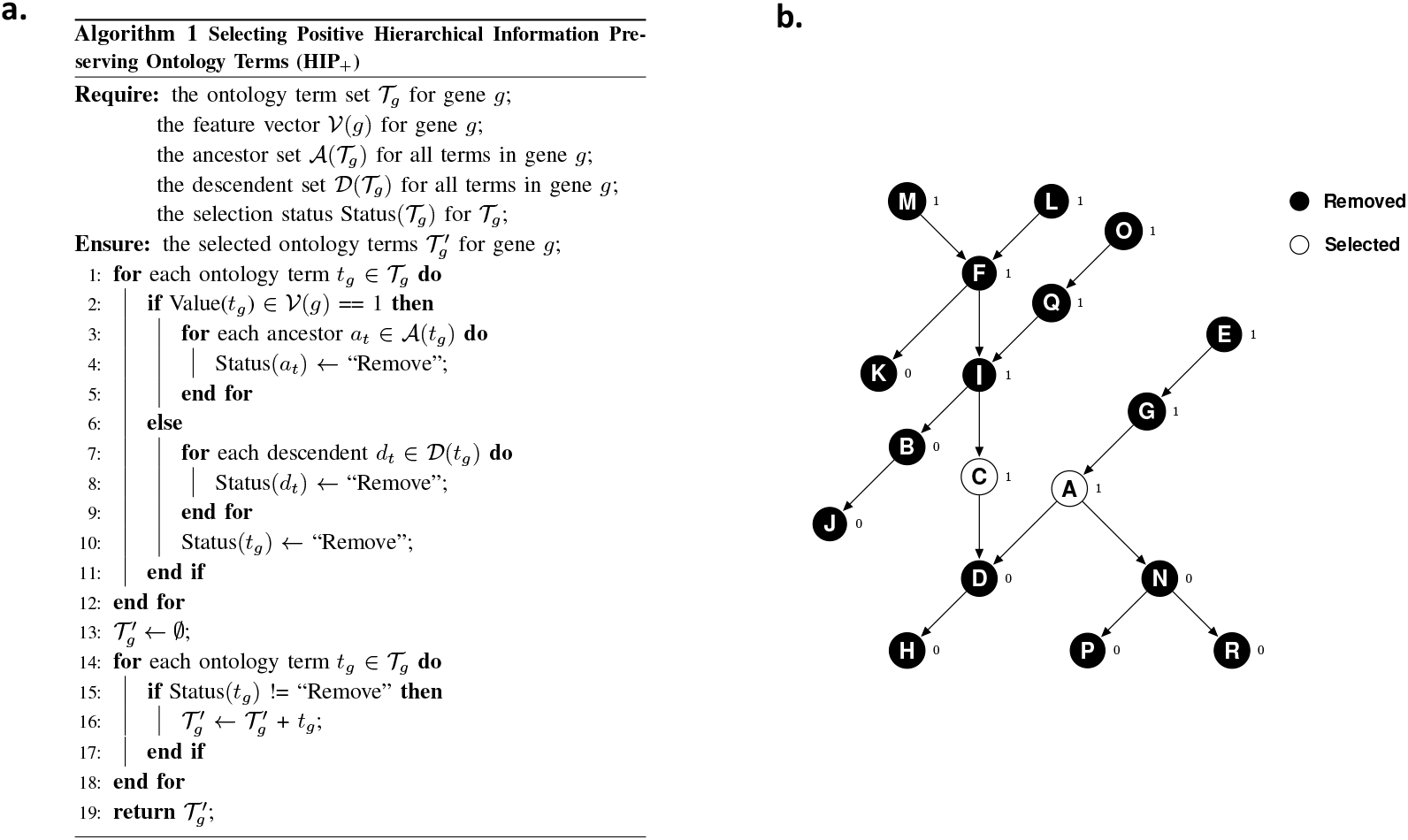
a. The pseudocode of the HIP_+_ hierarchical feature selection method. b. An example of the ontology terms selected by the HIP_+_ hierarchical feature selection method.

To explain how Algorithm 1 works, we use the example DAG shown in Figure 2.b, where 18 alphabets are used to denote 18 ontology terms. The binary values on the right side of the alphabet denote the annotation status, i.e. the value *1* denotes that the ontology term is annotated by a given gene, whereas the value *0* denotes that the ontology term is not annotated by a given gene. HIP_+_ conducts the hierarchical feature selection process for each individual term. For example, when processing the term C, as the value equals *1*, all term C’s ancestors (i.e. M, L, F, I, Q, and O) are removed, because all their feature values can be inferred by the feature value of term C. Analogously when processing term A, all its ancestor terms (i.e. G and E) are removed. Note that, HIP_+_ also removes all terms whose values equal *0*. For example, the terms K, B, J, D, H, N, P and R are removed. Finally, only terms C and A are kept, as denoted by the white background colour.

## 3 Results

We evaluated the predictive performance of the proposed HIP_+_ method by using different HPO label sets that are generated by different filtering threshold values *τ* and the HIP_+_ terms selection process. Figure 3.a shows the HPO terms distributions in different HPO terms label sets with corresponding *τ* values. The original HPO terms label set consists of 4,550 HPO terms for 184 ageing-related genes. After applying the HIP_+_ hierarchical feature selection process, 2,265 HPO terms that bear the most specific HPO definitions are kept in the label set. The number of remaining HPO terms decreased dramatically with the increased values of *τ*. For example, after applying a *τ* value of 10% and the HIP_+_ hierarchical feature selection process, the number of HPO terms decreased to 219. Analogously, the number of HPO terms further decreased to 95, if the *τ* Value increased to 20%.

**Figure 3.**
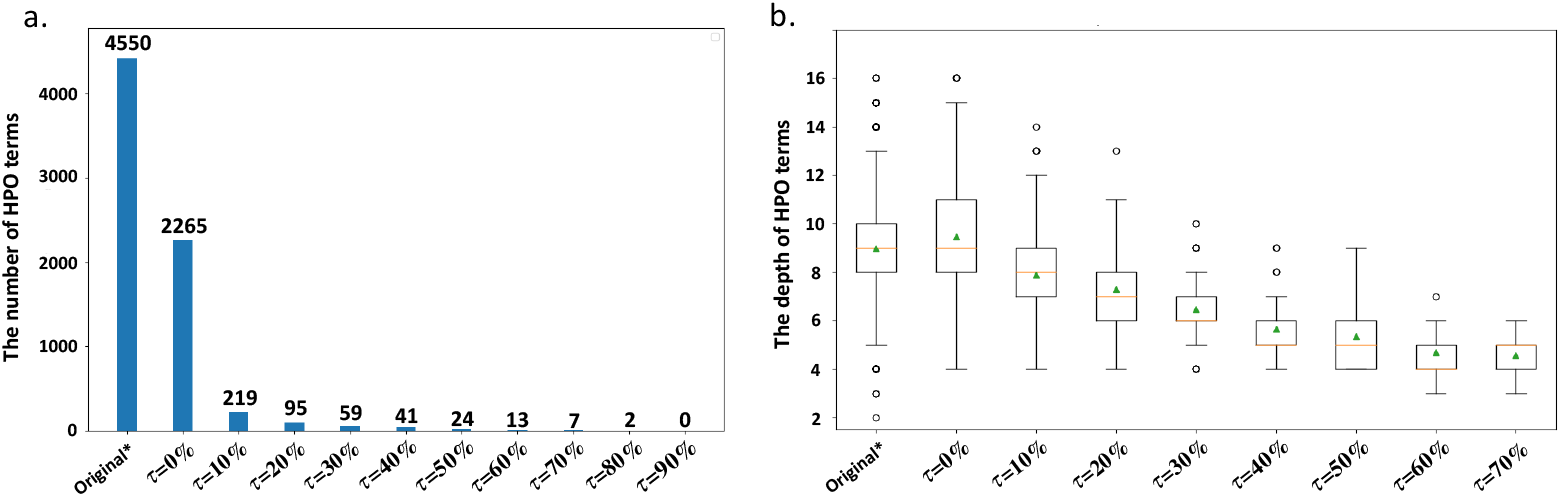
The charactistics of HPO terms in the dataset. a. The number of HPO terms in the class matrix after applying different thresholds. b. The median depth for HPO terms after applying different thresholds.

### 3.1 HIP_+_ successfully improved the accuracy of predicting HPO terms and outperformed the HIP method

We evaluated the predictive performance of the proposed HIP_+_ method to seven different HPO terms label sets which were created by using different filtering threshold values *τ* and the hierarchical feature selection process. In general, HIP_+_ obtained the highest predictive accuracy, compared with the conventional HIP method and using the entire GO terms feature set without feature selection. As shown in Table 1 (on the columns titled by *raw*), HIP_+_ obtained the highest median F1 scores and MCC values in 4 and 3 datasets, respectively. It also obtained the highest median average precision values in those two datasets. Conversely, the conventional HIP method failed to obtain any highest median F1 scores on those 7 HPO terms label sets – it only obtained the highest median MCC value and average precision score in 1 dataset, respectively. Analogously, another benchmark method, using the entire GO terms feature set without feature selection only obtained the highest median F1 scores and MCC values in two datasets, though it obtained the highest median average precision scores in 3 datasets.

**Table 1.** The performance of HIP_+_, HIP and the benchmark method for predicting different HPO terms label sets.

| Metrics | Feature types | $\tau=0\%$ | | $\tau=10\%$ | | $\tau=20\%$ | | $\tau=30\%$ | | $\tau=40\%$ | | $\tau=50\%$ | | $\tau=60\%$ | |
| --- | --- | --- | --- | --- | --- | --- | --- | --- | --- | --- | --- | --- | --- | --- | --- |
|  |  | Raw | Weighted | Raw | Weighted | Raw | Weighted | Raw | Weighted | Raw | Weighted | Raw | Weighted | Raw | Weighted |
| F1 | All | 0.000 | 0.000 | 0.050 | 0.120 | 0.164 | 0.239 | 0.367 | 0.341 | 0.493 | 0.350 | **0.656** | 0.373 | **0.801** | **0.349** |
|  | HIP | 0.000 | 0.000 | 0.067 | 0.180 | 0.235 | 0.292 | 0.397 | 0.344 | 0.522 | 0.344 | 0.581 | 0.348 | 0.725 | 0.344 |
|  | HIP<sub>+</sub> | 0.000 | 0.000 | **0.252** | **0.316** | **0.390** | **0.371** | **0.503** | **0.393** | **0.590** | **0.382** | 0.650 | **0.376** | 0.761 | 0.347 |
| MCC | All | 0.000 | 0.000 | 0.033 | 0.100 | 0.135 | 0.209 | **0.181** | 0.233 | 0.181 | 0.209 | **0.181** | **0.195** | 0.097 | 0.133 |
|  | HIP | 0.000 | 0.000 | 0.060 | 0.146 | 0.156 | 0.223 | 0.171 | **0.236** | 0.164 | **0.211** | 0.166 | 0.176 | **0.098** | **0.139** |
|  | HIP<sub>+</sub> | 0.000 | 0.000 | **0.156** | **0.245** | **0.164** | **0.238** | 0.163 | 0.220 | **0.186** | 0.203 | 0.159 | 0.167 | 0.080 | 0.123 |
| AP | All | 0.010 | 0.080 | 0.394 | **0.397** | **0.492** | **0.428** | **0.603** | 0.419 | **0.650** | **0.398** | 0.692 | 0.385 | 0.778 | 0.337 |
|  | HIP | 0.010 | 0.080 | 0.391 | 0.395 | 0.480 | 0.423 | 0.585 | 0.414 | 0.645 | **0.398** | **0.703** | **0.392** | 0.784 | **0.348** |
|  | HIP<sub>+</sub> | 0.010 | 0.080 | **0.403** | 0.395 | 0.487 | 0.420 | 0.576 | **0.424** | 0.638 | 0.387 | 0.685 | 0.388 | **0.799** | 0.344 |

### 3.2 HIP_+_ successfully improved the accuracy of predicting HPO terms that bear more specific human phenotypic abnormality definitions

As HPO terms are organized as a hierarchy, those terms located in the deeper position of the hierarchy bear more specific definitions, which are also more valuable in terms of biological knowledge discovery tasks, compared with those terms located in higher positions bearing more generic definitions. Also, due to the annotation quality issue, it is generally more difficult to make predictions on those terms bearing more specific definitions. As reflected by the results shown in Table 1, with the increased value of *τ*, the predictive accuracies for the corresponding HPO terms label sets increase too. For example, when *τ* equals 60%, only those terms bearing more generic definitions will be kept, as they received at least 110 genes annotations. Therefore, it is necessary to find the trade-off between predictive accuracy and interpretability for those different HPO terms label sets. We use the depth information for individual HPO terms to normalise the predictive accuracy for different HPO terms. As shown in Equations 1 and 2, the depth-weighted metric takes the square root of the product between the raw metric and the depth information. Here we use the longest distance to the root (i.e. # layer) as the measurement to calculate the depth for each individual HPO term.

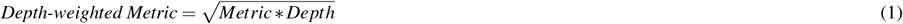

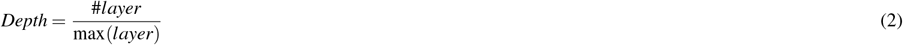

As shown in Table 1 (on the columns titled by *Weighted*), HIP_+_ obtained the highest median depth-weighted F1 scores in 5 out of 7 HPO terms label sets, and the overall highest depth-weighted F1 score (i.e. 0.393) with the *τ*=30% HPO terms label set.

In terms of MCC values, HIP_+_ obtained the highest median depth-weighted MCC values in 2 out of 7 HPO terms label sets, and the overall highest median depth-weighted MCC value (i.e. 0.245) with the *τ*=10% HPO terms label set. HIP didn’t obtain any highest median depth-weighted F1 score, but the GO terms selected by HIP obtained the highest median depth-weighted MCC values in 3 out of 7 datasets. The benchmark method that uses the entire GO terms feature set without feature selection only obtained the highest median depth-weighted F1 score and MCC value in 1 dataset, respectively. However, it obtained the highest median depth-weighted average precision value in 3 out of 7 datasets, and the overall highest median depth-weighted average precision value (i.e. 0.428) in the *τ*=20% HPO terms label set. Figure 4 shows the pairwise comparisons between HIP_+_, HIP and the benchmark method. It is obvious that HIP_+_ obtained higher depth-weighted F1 scores (Figure 4.a) and depth-weighted MCC values (Figure 4.d) in predicting the majority of HPO terms, compared with the benchmark method. The conventional HIP method also performed better than the benchmark method, as shown in Figure 4.b and Figure 4.e. HIP_+_ outperformed HIP – the former obtained higher depth-weighted F1 scores in 53 out of 59 HPO terms (Figure 4.c) and higher depth-weighted MCC values in 163 out of 219 HPO terms (Figure 4.f). However, both HIP_+_ and HIP failed to improve the depth-weighted average precision values, compared with the benchmark method, which obtained higher depth-weighted average precision values in more HPO terms, as shown in Figures 4.g and 4.h.

**Figure 4.**
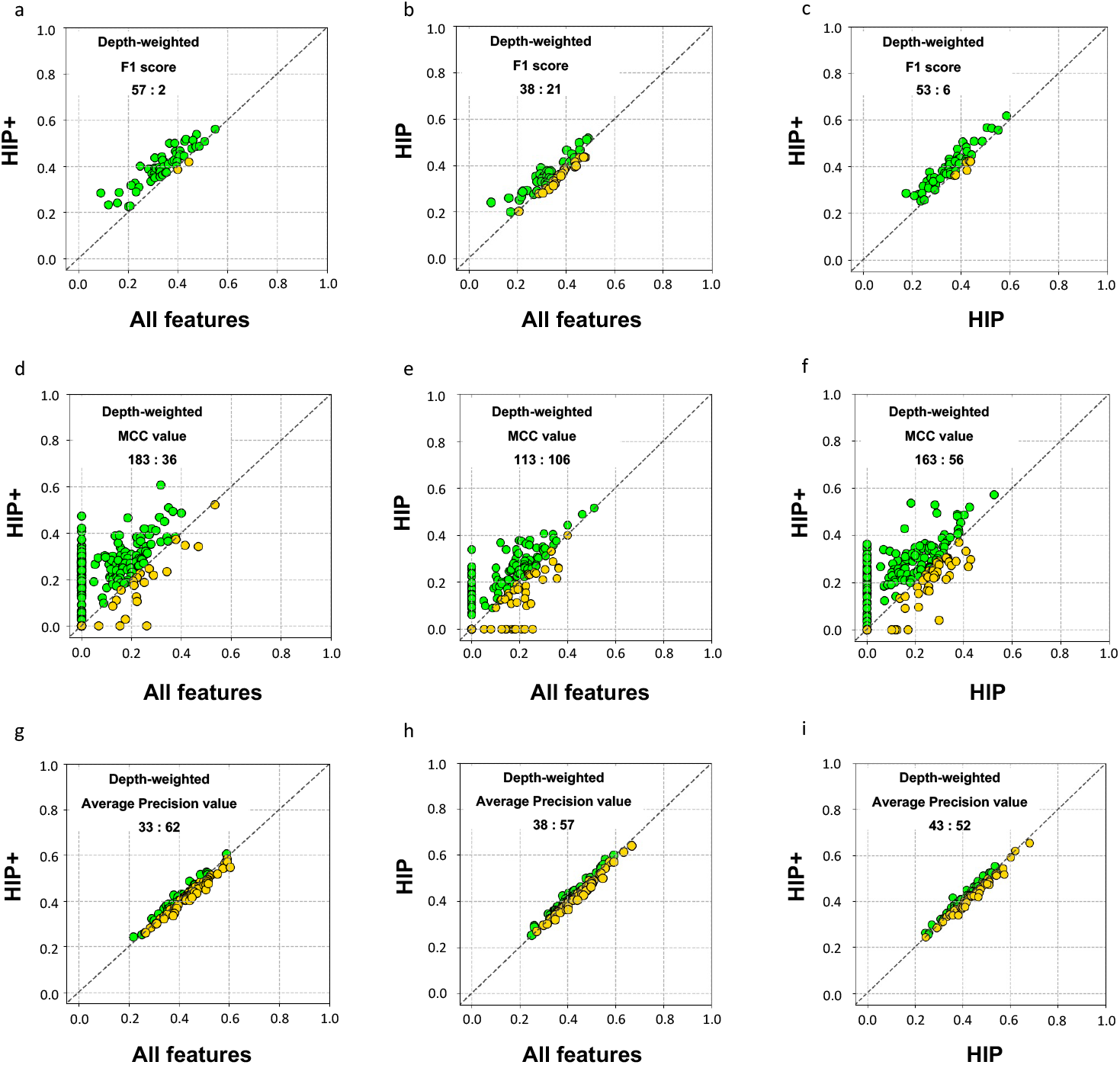
Comparison of depth-weighted F1 scores obtained by the *τ*=30% HPO terms label set (a-c), depth-weighted MCC values obtained by the *τ*=10% HPO terms label set (d-f) and depth-weighted average precision values obtained by the *τ*=20% HPO terms label set (g-i) obtained by HIP+, HIP and the benchmark method.

### 3.3 Biological interpretations on frequently selected GO terms

We discuss some important GO terms that highlight the relationships between ageing and human phenotypic abnormalities. We focus on the HPO label set generated by a *τ* value of 30% to make the discussion as it leads to the highest weighted F1 score. Table 2 displays four GO terms that were selected by HIP_+_ and bear high depth-weighted feature importance values, which are calculated by raw random forests feature importance (i.e. Gini importance) values and GO term hierarchy depth information (analogous to the weighted F1 score).

**Table 2.**
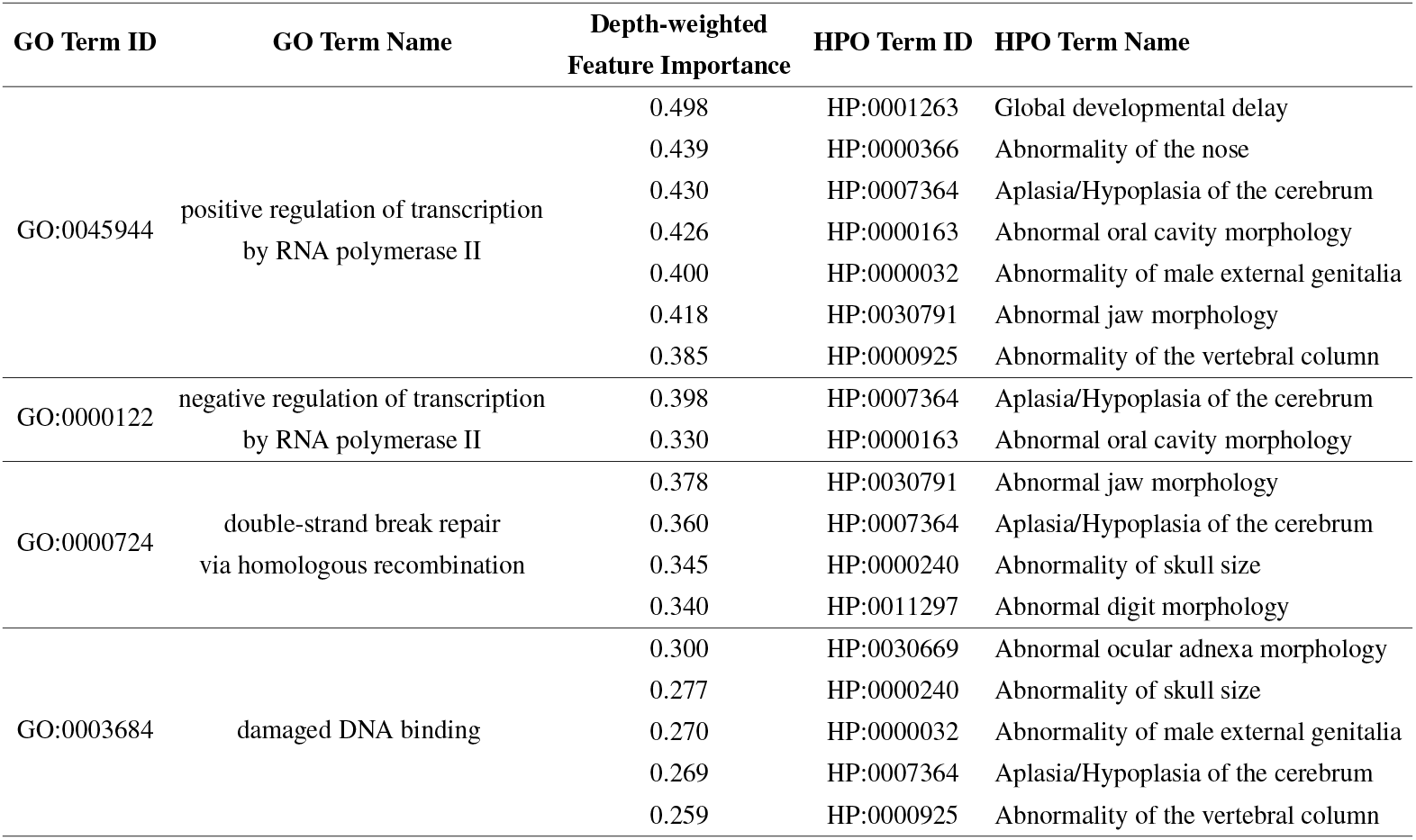
The GO terms selected by HIP_+_ with high depth-weighted feature importance values.

In general, all those four GO terms are related to transcription processes that play a crucial role in human ageing and development. It has been revealed that multiple ageing-realted transcription factors like DAF-16/FOXO^19–22^ are regulatory-related with pathways like insulin and insulin-like growth factor-1 (IGF-1)^23^ that are suppressed by DNA damage response^24^– a well-known ageing-related factor that causes cell dysfunction when associated DNA repair mechanism is flawed. It has also been found that some DNA repair pathways are associated with clinical phenotypes like Ataxia with oculomotor apraxia^1^ causing global developmental delay (HP:0001263) and Ataxia Telangiectasia^2^ – a type of Aplasia/Hypoplasia of the cerebrum (HP:0007364).

## 4 Conculsion

In this work, we proposed a novel hierarchical feature selection-based knowledge discovery framework to highlight some important associations between ageing processes and human phenotypic abnormalities. Future research could involve proposing other hierarchy-based feature selection or probabilistic graphical models to investigate novel dependencies between Gene Ontology and Human Phenotypic Abnormalities terms.

## Author contributions statement

C.W. conceived and conducted the experiments, C.W. and C.B. analysed the results. All authors reviewed the manuscript.

## Acknowledgement

We acknowledge the Birkbeck BEI School Impact Grant.

